# How symbiosis and ecological context influence the variable expression of transgenerational wing induction upon fungal infection of aphids

**DOI:** 10.1101/377077

**Authors:** Wen-Hao Tan, Miguel L. Reyes, Kim L. Hoang, Tarik Acevedo, Fredrick Leon, Joshua D. Barbosa, Nicole M. Gerardo

## Abstract

Aphids, like most animals, mount a diverse set of defenses against pathogens. For aphids, two of the best studied defenses are symbiont-conferred protection and transgenerational wing induction. Aphids can harbor bacterial symbionts that provide protection against pathogens, parasitoids and predators, as well as against other environmental stressors. In response to signals of danger, aphids also protect not themselves but their offspring by producing more winged than unwinged offspring as a way to ensure that their progeny may be able to escape deteriorating conditions. Such transgenerational wing induction has been studied most commonly as a response to overcrowding of host plants and presence of predators, but recent evidence suggests that pea aphids (*Acyrthosiphon pisum*) may also begin to produce a greater proportion of winged offspring when infected with fungal pathogens. Here, we explore this phenomenon further by asking how protective symbionts, pathogen dosage and environmental conditions influence this response. Overall, while we find some evidence that protective symbionts can modulate transgenerational wing induction in response to fungal pathogens, we observe that transgenerational wing induction in response to fungal infection is highly variable. That variability cannot be explained entirely by symbiont association, by pathogen load or by environmental stress, leaving the possibility that a complex interplay of genotypic and environmental factors may together influence this trait.

## Introduction

Animals have evolved several forms of defense against pathogens. Although the most studied defenses are mediated by cellular and humoral immune mechanisms, some defenses are mediated by behavioral mechanisms or symbiotic, microbial partners [1]. These alternative forms of defense may act in isolation or may influence one another.

Pea aphids *(Acyrthosiphon pisum)* utilize an array of defense strategies, and thus are a suitable system for studying how alternative forms of defense interact with one another. One defense, referred to here as transgenerational wing induction, arises from pea aphids’ reproductive biology. Under summer conditions, pea aphids asexually produce clonal copies of themselves, and, though genetically identical, these clonal offspring can be either unwinged (apterous) or winged (alate). For aphids, one commonly mounted defense against environmental stress, predators, and parasites is increased production of winged (relative to unwinged) offspring that can hopefully escape to better conditions [2–4]. Such transgenerational wing induction is similar to other transgenerational defenses, a phenomenon seen in many insect systems, in which defenses against pathogens are mounted not to protect oneself but to protect one’s offspring [5]. For example, immune-challenged bumblebees *(Bombus terrestris)* produce offspring with higher levels of antibacterial activity [6], and when infected with a protozoan parasite, monarch butterflies preferentially lay their eggs on plants that increase their offsprings’ resistance to the parasite [7]. Though aphid transgenerational wing induction is best known as a response to host plant overcrowding and presence of predators [3], recent work suggests that aphids utilize transgenerational wing induction in response to pathogen infection as well. Specifically, Hatano et al. [8] demonstrated that pea aphids can increase production of winged offspring in response to infection with the natural aphid pathogen, *Pandora neoaphidis.*

The finding that transgenerational wing induction is a potential aphid defense against fungal pathogens suggests that this form of defense could interact with another common aphid defense against fungal pathogens, namely association with protective bacterial symbionts. All pea aphids harbor an obligate endosymbiont, *Buchnera aphidicola,* that is essential for their survival and reproduction. In addition, individuals can harbor one to a few different facultative symbionts that can increase their hosts’ fitness [9–11]. Specifically, *Regiella insecticola,* a facultative endosymbiont, confers resistance against fungal pathogens [12], including *Pandora neoaphidis* [10], an important natural enemy in wild populations [13]. Interestingly, *R. insecticola* has been shown to impact transgenerational wing induction in response to crowding, suggesting the potential for the symbiosis to influence transgenerational wing induction in response to pathogens as well [14].

Our primary goal was to leverage the aphid - *Regiella* symbiont - *Pandora* pathogen system to explore how protective symbionts influence transgenerational defense. In our preliminary investigations, however, transgenerational wing induction in response to fungal infection was not consistently observed. To attempt to explain this variability, we also conducted a series of experiments to explore whether *R. insecticola* genotypes vary in their influence on transgenerational wing induction upon fungal infection, and whether the degree of pathogen exposure or environmental quality influences transgenerational wing induction upon fungal infection.

## Materials and methods

### Aphid lines

We used five lines of pea aphids *(Acyrthosiphon pisum*) previously established in the laboratory that have the same aphid genetic background but that harbor different genotypes of the secondary, facultative symbiont, *Regiella insecticola.* The five lines were established by experimentally infecting a single clonal aphid lineage, LSR1 [15], which did not have *Regiella,* with five genetically distinct *Regiella* (Ri, 313, 5.15, CO21, and U1), collected in previous studies [16–18], using established protocols [19, 20]. This created lines LSR1-Ri, LSR1-313, LSR1-5.15, LSR1-CO21, and LSR1-Ui, which we abbreviate here as LRi, L313, L515, LCO21, and LUi. In addition to these five lines, we also maintained a line without *Regiella* (LSR1-01, abbreviated as L01). Upon establishment, all aphid lines were reared asexually on fava beans *(Vicia faba)* in a temperature-controlled growth chamber at 20 °C under a 16/8h L/D cycle, which maintains them as asexual clones. Presence of symbionts was confirmed via PCR prior to conducting experiments [18, 21].

### *Pandora* fungal pathogen infections

*Pandora neoaphidis* ARSEF 2588 was obtained from the Agricultural Research Service Collection of Entomopathogenic Fungal Cultures, USA. We maintained *Pandora* in the laboratory by *in vivo* culturing, storing dead, infected aphids at 4 °C following methods described in Parker et al. [17]. We performed the fungal infection experiments using an established protocol [22] that mimics the natural route of pathogen transmission. Infected aphid cadavers, the fungal source, were placed on 1.5% tap water agar at 18 °C for 14-16 hours, providing sufficient time for the fungus to sporulate prior to aphid infections. Recently-molted (10-day old) adult aphids were experimentally infected by placing them in the bottom of an infection chamber (a PVC tube, 28 mm diameter and 40 mm height) on top of which we placed an agar plate with sporulating cadavers, allowing the experimental aphids to receive a fungal spore shower. Agar plates were rotated among infection chambers to homogenize the infection dosage, and a grid slide was used to estimate the infection dosage (number of spores / mm^2^). The infection period was 3-hr unless otherwise specified. Control aphids were handled similarly but were instead placed under agar plates without infected cadavers. After infection, we transferred aphids to two-week-old fava plants to monitor survival and offspring production. During the first four days post-infection, the plants were covered with solid plastic cups in order to keep the environment moist, as *Pandora* requires high humidity to infect aphids [23]. Afterward, the plants were covered by plastic cups with mesh tops.

### Overview of survival and wing induction measurements

We used survivorship to quantify the differences in *Pandora* resistance between aphid lines and measured induction of winged offspring production as a transgenerational defense trait. For survival assays, we inspected infected and uninfected aphids daily to record survival. Dead aphids were checked for visible signs of sporulation. We monitored survival for 9-10 days, as infection-caused mortality and sporulation usually occur between 4-10 days after exposure in this system [22]. For transgenerational wing induction, we collected offspring produced in the four days post fungal infection by transferring each adult aphid to a new plant every other day. We recorded the number of offspring produced each day. The proportion of offspring that were winged was recorded after each cohort reached adulthood.

### Experiment 1: Influence of *Regiella* presence on transgenerational wing induction upon *Pandora* infection

We used two aphid lines, LRi (harboring *Regiella)* and L01 (without *Regiella).* We exposed 34 aphids of each line to *Pandora*, and monitored 34 control, uninfected aphids per line as well. For each treatment group, 10 aphids went to individual plants to monitor offspring production, and 24 aphids were monitored (8 aphids on each of three plants) for survival. We monitored survival of the exposed (F0) aphids and assessed the proportion of their offspring (F1) that were winged using methods detailed above. After the F1 aphids became adults, we typically randomly selected six unwinged, F1 offspring of each F0 individual and transferred them to individual plants to monitor the winged status of the F2 generation. Twenty-two F0 individuals produced fewer than six unwinged offspring however, and we thus used fewer offspring from these individuals (F1 per F0: range = 1-6, median = 5).

### Experiment 2: Influence of alternative *Regiella* lines on transgenerational wing induction upon *Pandora* infection

Given that we did not observe wing induction in either *Regiella*-present or Regiella-absent lines in Experiment 1 in response to fungal infection, and that Regiella-mediated resistance against *Pandora* is dependent on genotype-by-genotype specificity [17], we hypothesized that *Regiella’s* influence on transgenerational wing induction could also be genotype-specific. In this experiment, we tested for the effect of symbiont genotype on resistance and wing induction upon fungal infection. We used all six lines of aphids described above (five lines harboring different genotypes of *Regiella* and one without *Regiella).* We performed the experiment twice. We first conducted the experiment using previously established lines, and then repeated this experiment with re-established lines to ensure that the host genetic background was identical across all lines (while aphids clonally reproduce, mutations can occur and become fixed in lab lines). We conducted infections and monitored survival of F0 individuals, and measured the proportion of their F1 offspring that were winged following methods described above, with the exception that in the first experiment we did a 3-day rather than 4-day collection of F1 offspring due to logistical constraints. Although the same protocol and infection period was used, the fungal dosages were very different between the two replicates. The fungal infection dosage was 12.3 spores/mm^2^ in Replicate A and 105.6 spores/mm^2^ in Replicate B. For Replicate A, sample sizes ranged from 3 – 10 (median = 6.5) per treatment group; for Replicate B, sample sizes ranged from 11 – 12 (median = 12) per treatment group. For Replicate B, aphid line L313 was removed from analyses due to low survival of the control group (all died within 10 days).

### Experiment 3: Influence of pathogen dose on transgenerational wing induction upon *Pandora* infection

Given that the two replicates of Experiment 2 showed different results in terms of transgenerational offspring production in response to *Pandora* infection, we attempted to examine the potential factors influencing this response. A previous study demonstrated that higher infection dosage leads to higher pathogen burden and higher mortality in this system [22]. Due to the fact that the infection dosage was markedly different in the two replicates of Experiment 2, we hypothesized that induction of winged offspring production might be dependent on pathogen load. To test this, we exposed two aphid lines (LRi and L01) to three infection dosages: high (144.1 spores/mm^2^), medium (12.5 spore s/mm^2^), and low (1.6 spores/mm^2^) by altering the infection period to manipulate infection dosage. That is, a higher dosage group was infected for a longer time. In order to control for a confounding effect that staying longer in a chamber could increase stress, we added an uninfected control group for each infection period. Thus, this experiment was a fully factorial design of three infection periods (which correlates with dosage), two infection treatments (infected and uninfected), and two aphid lines. Sample sizes ranged from 10 – 12 (median = 11) per treatment group. After fungal infection, we followed the methods described above to monitor survival and offspring production.

### Experiment 4: Influence of environmental condition on transgenerational wing induction upon *Pandora* infection

Our results in Experiment 3 did not detect increased production of winged offspring under high pathogen load, suggesting that pathogen dosage alone is not the main factor triggering the expression of this response. Through comparing conditions in the above experiments, we hypothesized that other factors, such as host plant quality and aphid health, could influence the production of winged offspring. To manipulate condition, in Experiment 4, we used starvation and drought as abiotic stresses, both of which have been shown to negatively affect pea aphids [24–26]. For the starvation treatment, prior to fungal infections, we starved aphids for 12 hours when they were four days old (young nymph) and again when they were 10 days old (newly-molted adults). To do so, we moved aphids that were reared on the same plant to another pot with moist soil but no plants. Aphids were transferred back to their original plant after the starvation treatment. For drought treatments, we transplanted fava plants to dry soil the same day we transferred aphids onto them. Those two treatments caused the aphids to look pale, which is an indication of poor condition [27]. This experiment was thus a fully factorial design of three environmental conditions (starvation, drought, and control), two infection treatments (infected, uninfected) and two lines (LRi and L01). We used seven aphids for each treatment group. We followed the same infection protocol as above, using an infection period of eight hours to reach a medium dosage (21.0 spores/mm^2^). After fungal infection, we assayed survival and offspring production as above.

### Statistical analyses

Across all experiments, we measured survivorship and the proportion offspring that were winged and tested for the effects of several factors. Survival analyses were performed using Cox Proportional Hazardous models using the R package survival 2.41-3 [28]. The proportions of offspring that were winged were analyzed using General Linear Models (GLMs) with either binomial or quasi-binomial error structures, depending on the dispersion parameter. All analyses were performed using R version 3.4.1 [29].

## Results

### Experiment 1: Influence of *Regiella* presence on transgenerational wing induction upon *Pandora* infection

Fungal infection significantly reduced aphid survival, and aphid lines differed in their survival; however, there was no significant interaction between infection status and aphid line (Fig 1A; infection: χ^2^ = 8.198, df = 1, *P* = 0.004; line: χ^2^ = 14.938, df = 1, *P* < 0.001; interactions: χ^2^ = 0.270, df = 1, *P* = 0.604). *Pandora* fungal infection of F0 generation aphids did not induce production of winged offspring relative to control, uninfected aphids (Table 1). Specifically, while some F1 offspring were winged, and symbiont status significantly affected this trait, it was not influenced by fungal infection of their mothers (Table 1; Fig 1B). The F2 generation consisted of few winged: two out of 2854 F2 offspring were winged. Specifically, out of a total of 195 F1 aphids, only two of them produced a winged offspring.

**Table 1.**
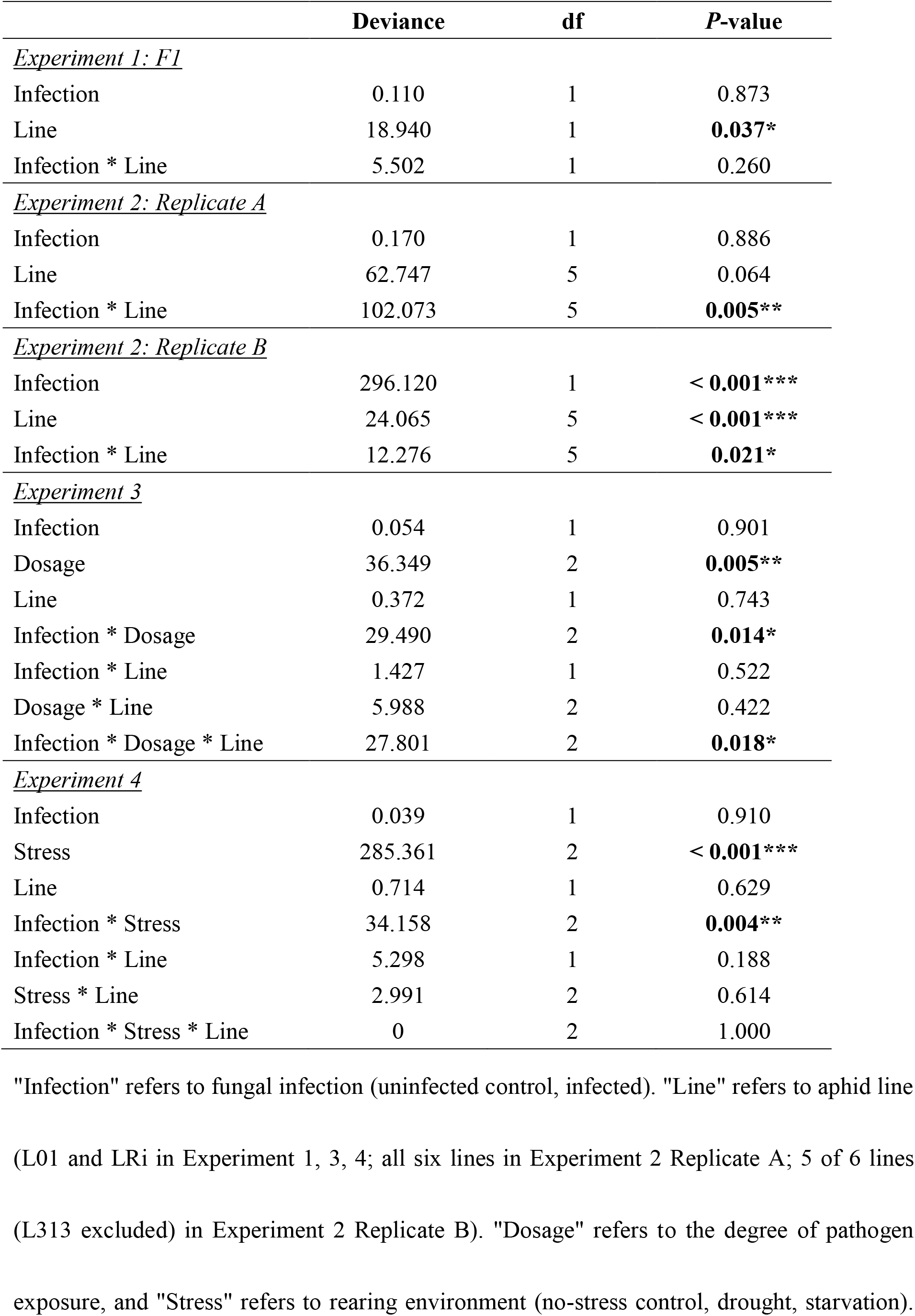
Results of analysis of deviance of GLMs for the proportion of offspring that were winged for the four experiments.

**Fig 1.**
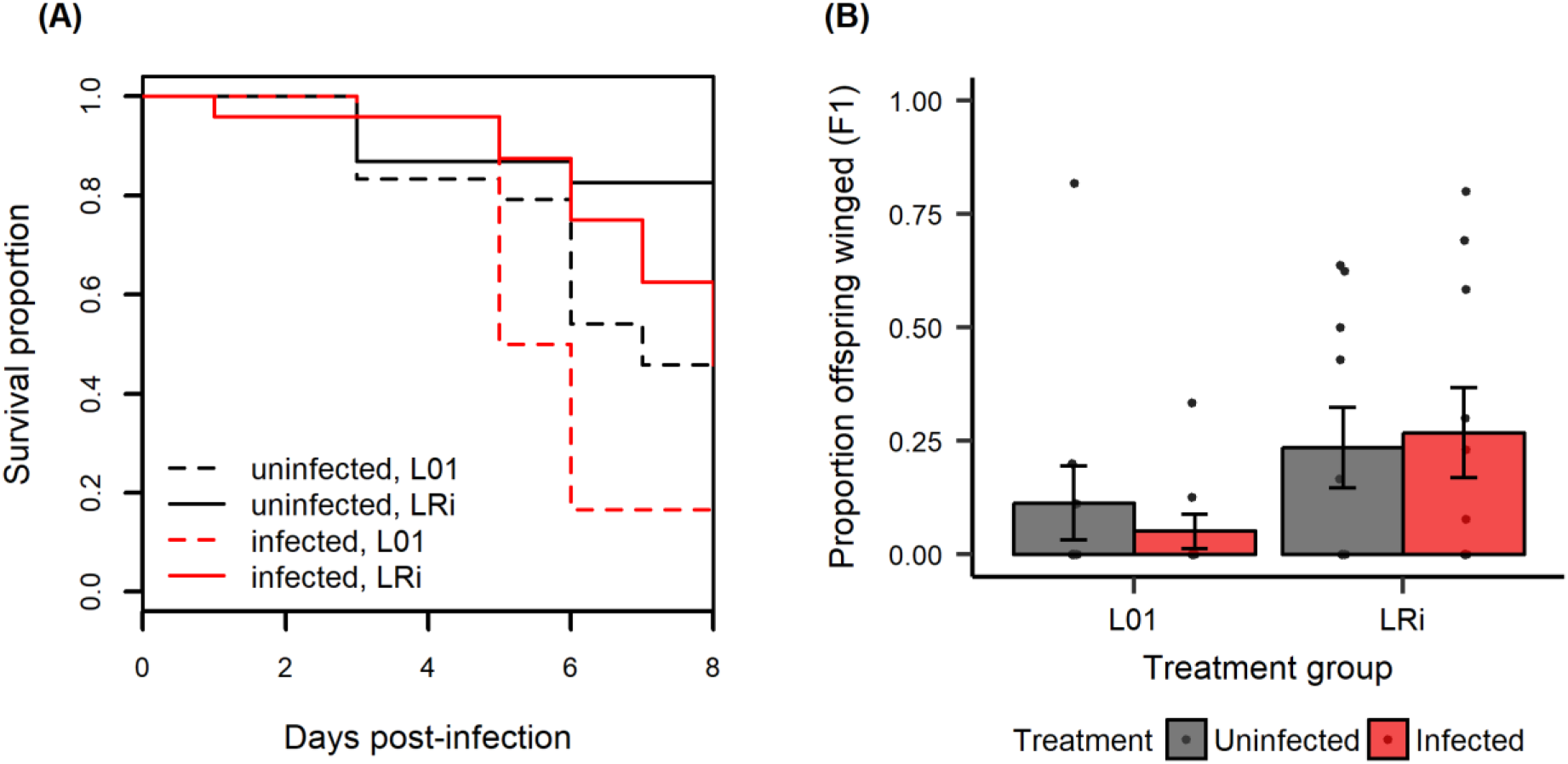
Experiment 1. Fungal infection reduced survival but had no effect on the proportion of offspring that were winged across two aphid lines. F1 are offspring of experimentally infected or control F0 mothers. (A) F0 aphid survival upon *Pandora* infection. Solid lines indicate aphid line LRi (with *Regiella)* and dotted lines indicate line L01 (without *Regiella);* (B) the proportion of F1 offspring that were winged. An average of 10.3 offspring were produced per F0 aphid over four days. Sample sizes range from 9 - 10 (median = 10) F0 monitored for winged offspring production per treatment group. Points are proportion of winged offspring for each F0 individual; bars represent mean ±1 SEM.

In Experiment 1, F1 are offspring of experimentally infected F0 mothers. While we also collected data on the F2 offspring of F1 aphids, the results for F2 were not analyzed because very few F2 winged offspring were produced: out of a total of 195 F1 aphids used, only two of them produced a winged offspring (2 out of 2854 offspring in total).

### Experiment 2: Influence of alternative *Regiella* lines on transgenerational wing induction upon *Pandora* infection

#### Replicate A

Fungal infection significantly reduced aphid survival (Fig 2A; χ^2^ = 64.907, df = 1, *P* < 0. 001), and symbiont genotype had a significant effect on host survival upon fungal infection (χ^2^ = 29.908, df = 5, *P* < 0.001). Resistance against *Pandora* differed between the six aphid-lines: LRi, LUi, and L313 had significantly higher survival than L01, while L515 and LCO21 did not (Fig 2B, Table 2). Neither fungal infection nor aphid line had significant main effects on the proportion of offspring that were winged; however, the interaction term had a significant effect, suggesting a role for symbionts in altering induction of winged offspring production (Fig 2C, Table 1).

**Table 2.**
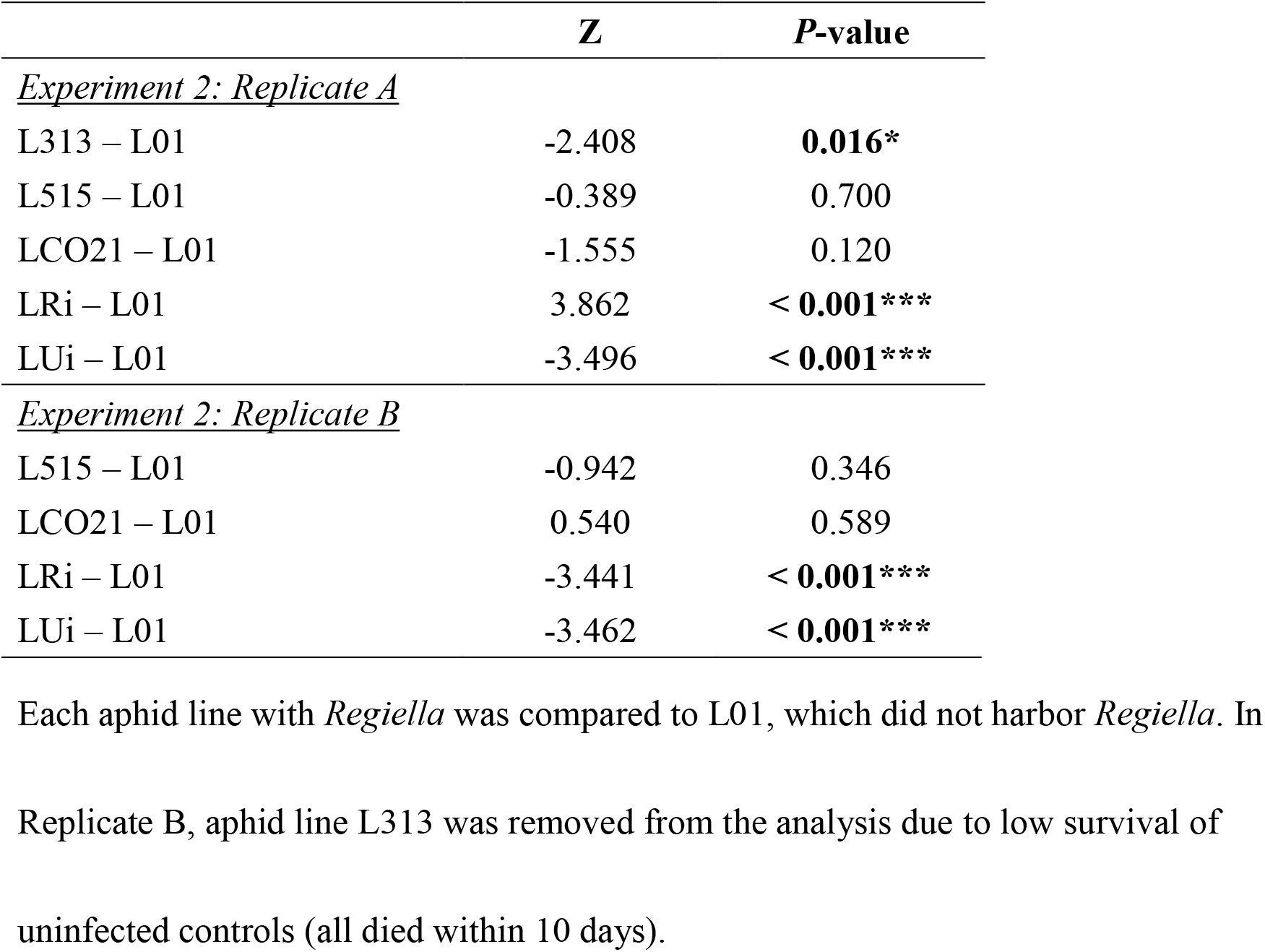
Differences in survival between aphid lines in Experiment 2.

**Fig 2.**
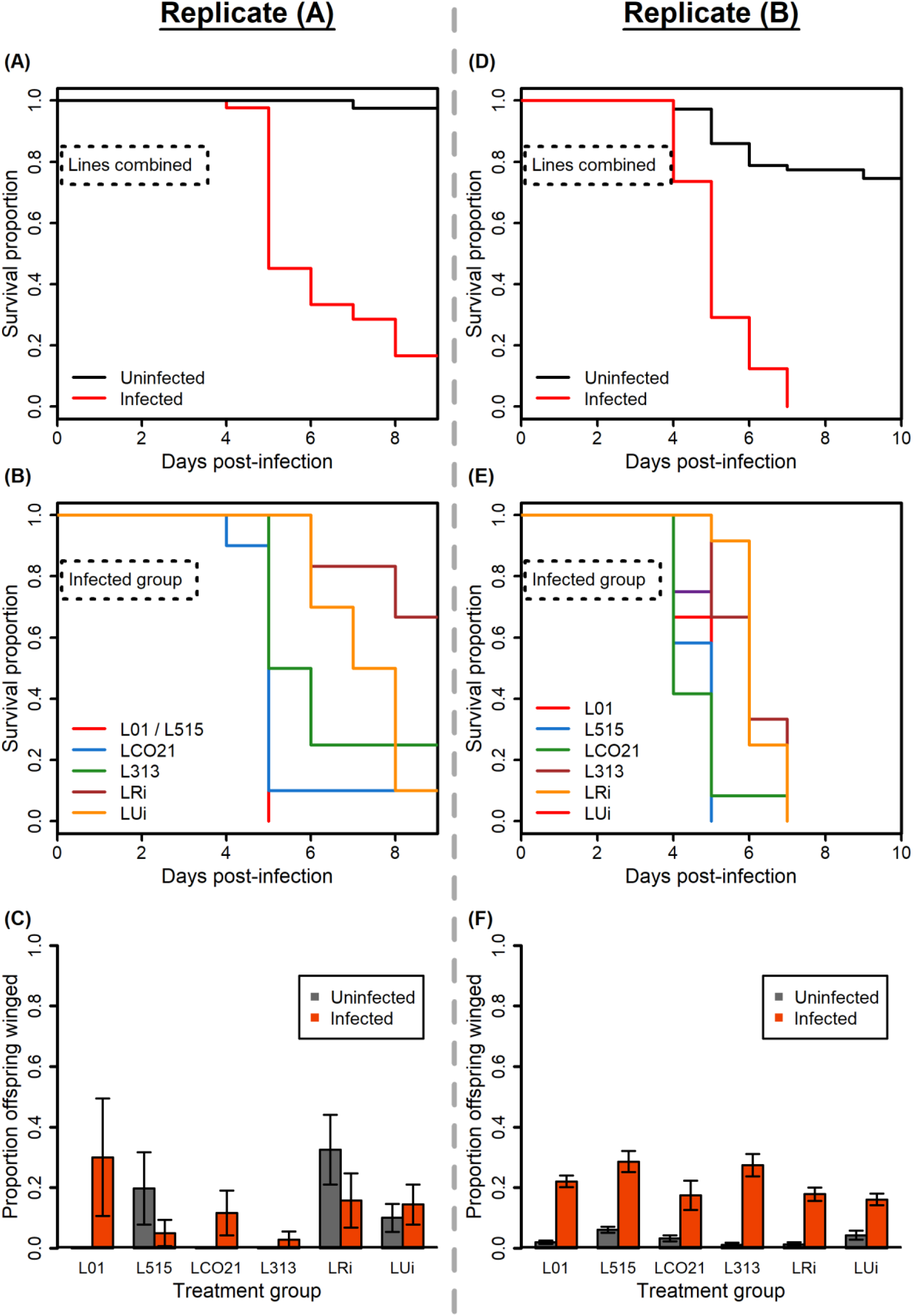
Experiment 2. Lines with different symbionts varied in terms of survival upon fungal infection but did not consistently vary in terms of winged offspring production. Replicate A shown in (A) – (C), and Replicate B shown in (D) – (F). **(A) and (D):** survival upon *Pandora* infection relative to control, uninfected aphids - aphids lines combined. In both replicates, fungal infection reduced aphid survival. **(B) and (E):** survival upon *Pandora* infection – aphid lines plotted separately for pathogen infected groups only. In both replicates, *Regiella* genotype impacts aphid survival upon infection. Note that in (B), the survival curve for L01 and L515 are shown as the same color because the curves completely overlap. **(C) and (F):** the proportion of offspring that were winged upon fungal infection. In Replicate A, an average of 22.1 offspring per experimental aphid were produced across three days; in Replicate B, an average of 25.2 offspring were produced across four days. In Replicate A, fungal infection did not consistently induce an increase in winged offspring production; however, in Replicate B, fungal infection consistently induced an increase in winged offspring production. Although the two replicates showed strikingly different patterns, there was a significant interaction between infection and aphid line found in both replicates, suggesting that symbiont genotypes alter responses to fungal infections in different ways. Error bars in (C) and (F) represent mean ±1 SEM. For Replicate A, sample sizes range from 3 – 10 (median = 6.5) experimental aphids monitored for survival and proportion of winged offspring produced per treatment group. For Replicate B, sample sizes range from 11 – 12 (median = 12) experimental aphids monitored for survival and proportion of winged offspring produced per treatment group.

#### Replicate B

Consistent with Replicate A, fungal infection significantly reduced aphid survival (Fig 2D; χ^2^ = 115.301, df = 1, *P* < 0.001), and symbiont genotype had a significant effect on survival upon fungal infection (χ^2^ = 26.099, df = 4, *P* < 0.001). Resistance against *Pandora* differed between aphid lines: LRi and LUi had significantly higher survival than L01, while L515 and LCO21 did not (Fig 2E, Table 2). Though patterns of survival were similar to Replicate A, transgenerational wing induction was strikingly different. The proportion of offspring that were winged was significantly influenced by fungal infection, aphid line, and their interaction (Fig 2F, Table 1).

### Experiment 3: Influence of pathogen dose on transgenerational wing induction upon *Pandora* infection

Fungal infection, infection dosage, and aphid line all had significant effects on host survival upon fungal infection (Fig 3; infection: χ^2^ = 44.542, df = 1, *P* < 0.001; dosage: χ^2^ = 13.378, df = 1, *P* < 0.001; line: χ^2^ = 12.264, df = 1, *P* < 0.001); higher dosages caused higher mortality in both lines. Fungal dosage had a significant effect on the proportion of offspring that were winged, as did the interaction between dosage and infection and the three-way interaction between infection, dosage, and aphid line (Fig 4, Table 1). However, the proportion of offspring that were winged was generally low: across all fungally infected aphid treatments only 46 of 1666 offspring (2.76%) were winged.

**Fig 3.**
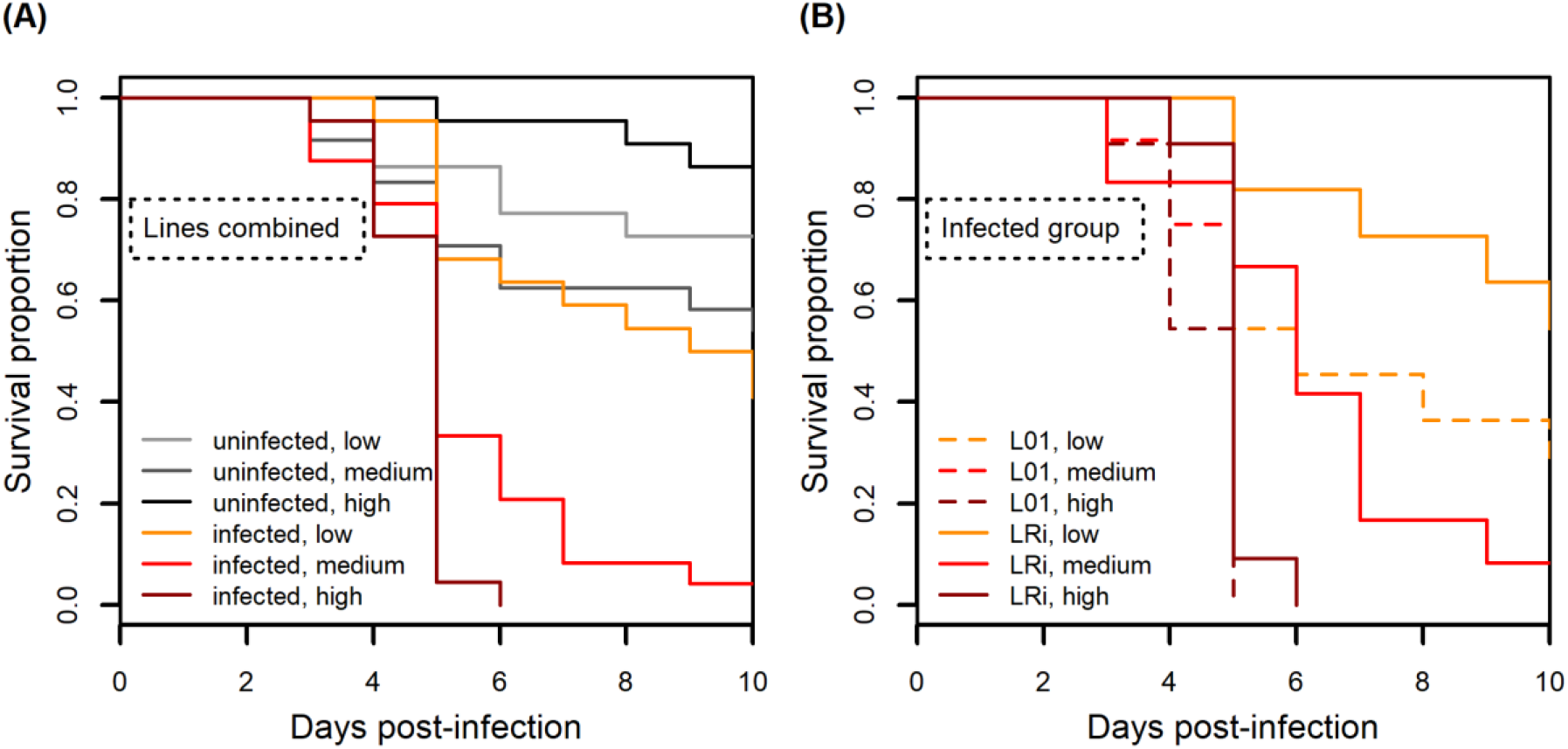
Experiment 3. Higher fungal dosage led to lower survival upon infection across two aphid lines. Each infection dosage has a corresponding control group in order to control for the effect of infection period (*i.e.*, the time aphids stayed in infection chambers). Low dose represents 1.6 spores/mm^2^, medium dose represents 12.5 spores/mm^2^, and high dose represents 144.1 spores/mm^2^. (A) aphid lines combined; (B) aphid lines plotted separately for infected groups only. Solid lines indicate aphid line LRi (with *Regiella),* and dotted lines indicate line L01 (without *Regiella).* Sample sizes range from 10 – 12 (median =11) per treatment group.

**Fig 4.**
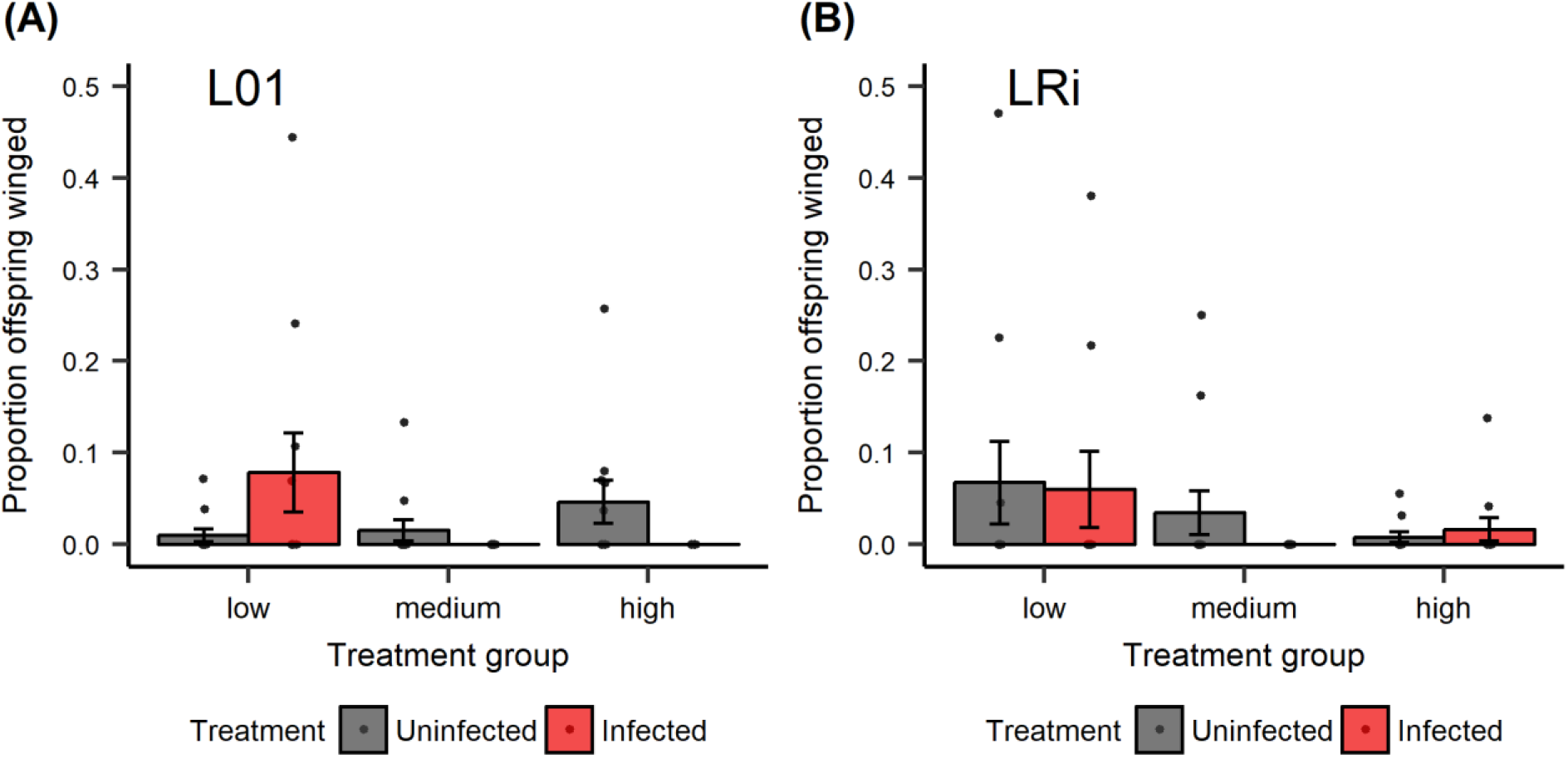
Experiment 3. The effects of infection, fungal dosage, and aphid lines on the proportion of offspring that were winged. Each infection dosage has a corresponding control group in order to control for the effect of infection period *(i.e.,* the time aphids stayed in infection chambers). Low dose represents 1.6 spores/mm^2^, medium dose represents 12.5 spores/mm^2^, and high dose represents 144.1 spores/mm^2^. (A) aphid line L01 (B) aphid line LRi. In this experiment, the proportion of offspring that were winged was generally low: across all fungally infected aphid treatments only 46 of 1666 offspring (2.76%) were winged (Note that the y-axis scale is 0 to 0.6). An average of 24.4 offspring were produced per experimental aphid across four days. Sample sizes range from 10 – 12 (median =11) experimental aphids monitored for proportion of winged offspring produced per treatment group. Points represent proportion of offspring that were winged for a particular individual; bars represent mean ±1 SEM.

### Experiment 4: Influence of environmental condition on transgenerational wing induction upon *Pandora* infection

A high background death rate was observed across treatments in this experiment. Neither the main effects of infection, environmental condition, nor aphid line had significant effects on host survival upon infection (Fig 5; infection: χ^2^ = 0.001, df = 1, *P =* 0.982; stress: χ^2^ = 0.437, df = 1, *P =* 0.804; line: χ^2^ = 0.695, df = 1, *P =* 0.404). Environmental condition, however, had a significant effect on the proportion of offspring that were winged, as did the interaction between stress and infection (Fig 6, Table 1). The most striking influence on winged offspring was starvation of mothers, which stimulated transgenerational winged offspring production in both the presence and absence of fungal infection.

**Fig 5.**
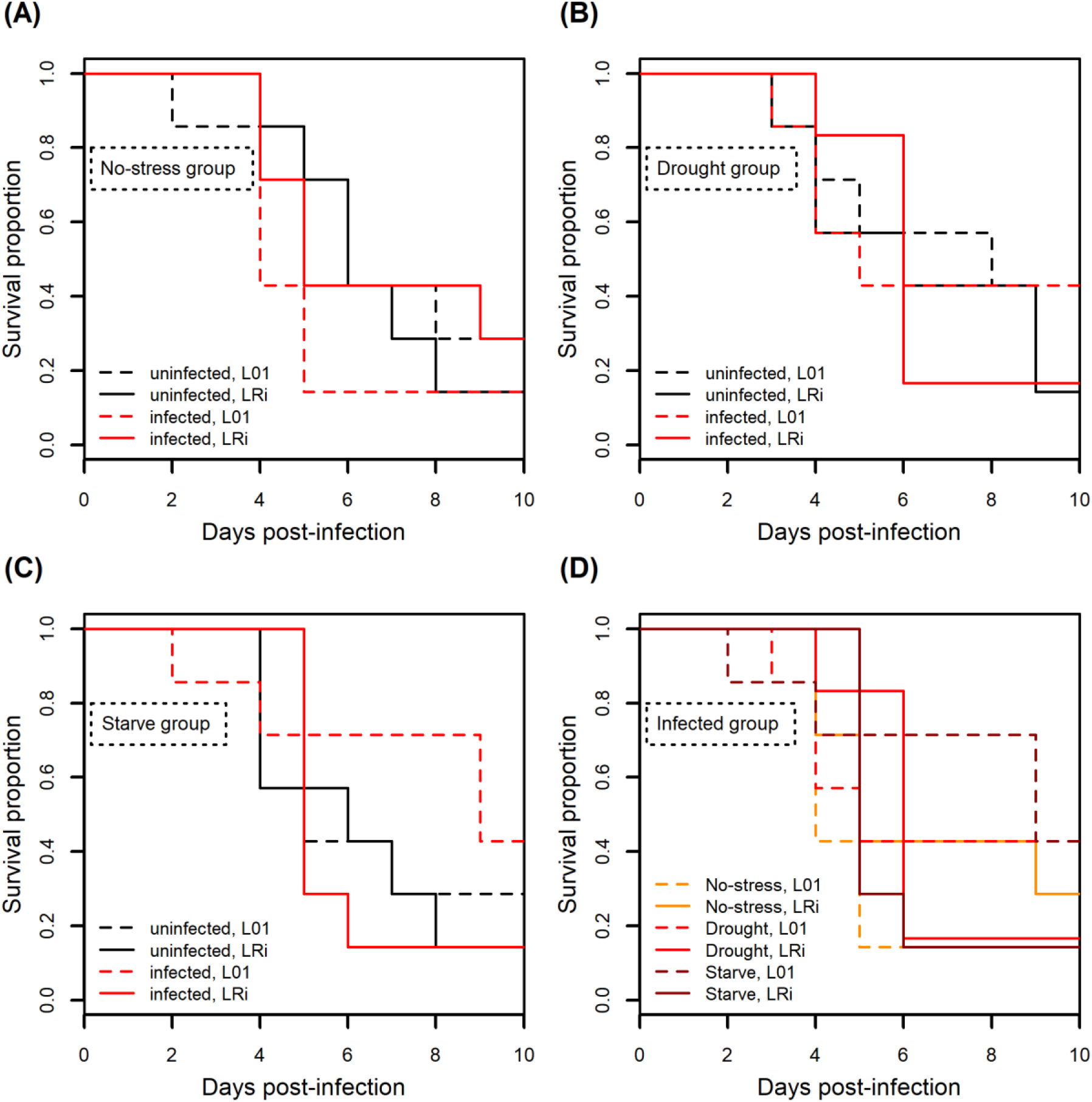
Experiment 4. Stress, infection, and aphid line had no effect on aphid survival upon fungal infection. Solid lines indicate aphid line LRi (with *Regiella),* and dotted lines indicate aphid line L01 (without *Regiella).* (A) no-stress (control) treatment; (B) drought treatment; (C) starvation treatment; (D) survival of infected aphids only, all treatments. Sample sizes equal to seven individuals per treatment group.

**Fig 6.**
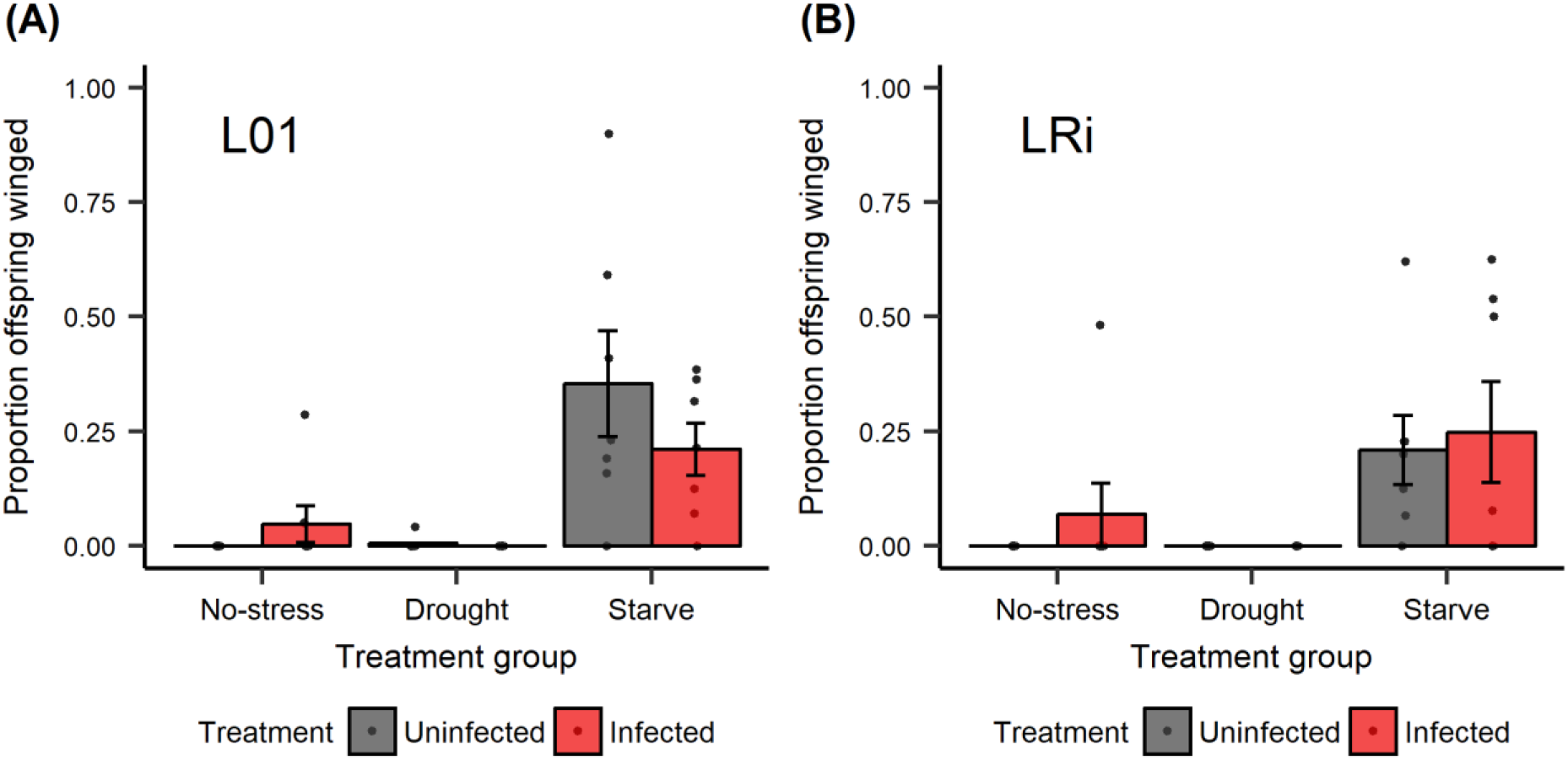
Experiment 4. Starvation induced an increase in the production of offspring that were winged but fungal infection did not. (A) aphid line L01, and (B) aphid line LRi. An average of 21.2 offspring were produced per experimental aphid across four days. Sample sizes equal to seven experimental aphids per treatment group. Points represent proportion of offspring that were winged for a particular individual; bars represent mean ±1 SEM.

## Discussion

The utilization and efficacy of defenses is often dependent on genotype-by-genotype interactions and on environmental context [30, 31]. In this study, we tested for the influence of multi-generational effects, symbiont genotype, pathogen dosage, and two abiotic stressors on the expression of an induced defense trait – production of winged offspring in response to fungal infection. Though transgenerational wing induction in response to fungal infection was reported in a previous study [8], we did not consistently observe wing induction. Our results suggest that wing induction of pea aphids upon *Pandora* infection may be strongly dependent on particular factors that were not captured in our experimental design. Given that the aphid and *Pandora* genotypes we used were different from the ones used in Hatano et al. [8], one possibility is that the response may vary between host genotypes, may vary based on pathogen genotypes, or may be influenced by genotype-by-genotype interactions. However, in our study, replicate experiments *(e.g.,* Experiment 2 Replicates A and B) exhibited considerable differences, suggesting that environmental variation not captured here also influences expression of this defense.

In Experiment 1, we tested whether exposure to pathogens leads to production of more winged daughters and/or more winged granddaughters. Our consideration of the potential influence on granddaughter physiology stems from the interesting reproductive biology of aphids. In days with relatively high levels of light, pea aphids reproduce via parthenogenesis, producing offspring as first instar nymphs (viviparous). Prior to birth, their developing embryos already have embryos developing within them, a condition known as telescoping generations [32]. As a result, it creates an opportunity for aphids to receive maternal and even grandmaternal signals related to environmental conditions and pathogen exposure [33]. Because of previous findings that aphids produced a greater proportion of winged offspring in response to fungal infection [8], we asked whether this pathogen exposure could have longer lasting effects across multiple generations. As we saw no influence of pathogen exposure on winged offspring production in either generation, this is not a consistently observed phenomenon. However, future work should test this interesting hypothesis in relation to other pea aphid defense traits.

Given that we saw little evidence of transgenerational wing induction in response to fungal infection in Experiment 1, we carried out a set of experiments to explore how genetic and non-genetic (environmental) variation might influence this trait. In Experiment 2, we asked whether symbiont genotypes varied in their effects on transgenerational wing induction in response to fungal infection. This built on previous research showing that *R. insecticola* genotypes vary in the level of protection that they confer against *P. neoaphidis* [17]. In the first replicate of the experiment, we saw no significant main effect of either fungal infection or symbiont genotype on the production of winged offspring, but the interaction between infection and symbiont genotype was significant, suggesting that lines with alternative symbionts varied in their response to infection. In contrast, in the second replicate, all aphid lines, regardless of symbiont genotype, produced more winged offspring when fungally infected, though the strength of the response varied across lines. Despite the inconsistent main effects of fungal infection and symbiont genotype, the fact that both experimental replicates demonstrated significant symbiont genotype by fungal infection interactions suggests that *Regiella* may play a role in regulating the response and that this regulation is likely symbiont genotype-specific.

Given the differences observed between the outcomes of the two replicates of Experiment 2, we asked what environmental conditions could have differed between the two replicates as a way to begin to understand the variable expression of transgenerational wing induction in response to fungal infection. We first noted that fungal pathogen virulence was higher in Replicate B than in Replicate A (Fig 2 (A) and (D)), consistent with the fact that fungal dosage in Replicate B (105.6 spores/mm^2^) was much higher than that in Replicate A (12.25 spores/mm^2^). We thus hypothesized that pathogen virulence and/or dosage could influence the expression of the defense trait. Results of Experiment 3 showed that higher fungal dosages led to lower survival, which is consistent with previous studies [22]. However, higher dosage did not result in higher proportions of winged offspring. Indeed, we instead observed fewer winged offspring produced when aphids were exposed to a higher pathogen dose. A recent study suggested that wing polyphenism in pea aphids is controlled by the ecdysone pathway – downregulation of the ecdysone pathway leads to increased winged offspring and vice versa [34]. Ecdysone is also identified as a positive regulator of innate immune mechanisms in other systems [35]. Therefore, it is possible that enhanced immune responses in the face of stronger pathogen challenge could lead to suppression of wing induction; however, future studies are required to disentangle the physiological mechanisms underlying this response.

Host defenses are often dependent on environmental-context [36]. For example, in honey bees, birds and humans decreasing nutrient availability decreases immunocompetence [37–39]. We hypothesized that the variation that we observed between the two replicates of Experiment 2 could be a result of environment variation that influenced host condition. Specifically, control, uninfected aphids had lower survival in Experiment 2 Replicate B compared to Replicate A, suggesting that their overall condition may have been worse. In Experiment 4, we attempted to modulate host condition by rearing aphids under control, drought and starvation conditions. While starvation triggered a strong induction of winged offspring production, neither starvation nor drought enhanced responses to fungal infection. We should note, however, that fungal infection did not significantly impact aphid survival in this experiment, though we did observe sporulating aphids suggesting that the aphids were indeed infected. Thus, it is possible environmental stress could enhance transgenerational winged offspring production in response to fungal pathogen under different infection conditions.

## Conclusions

In this study, through a series of experiments that tested the influence of multiple factors on transgenerational wing induction in response to *Pandora* infection, we did not consistently observe the increased production of winged offspring by infected individuals. Our results confirmed that *Regiella* genotypes differ in the strength of protection that they confer to aphids, and showed that wing induction, though not consistently expressed, may be dependent on symbiont genotype to some extent. Our study further suggested that *Pandora*-induced winged offspring production may be strongly dependent on other environmental or non-genetic factors not captured in our experiments, and may have strong specificity across host, symbiont, and pathogen genotypes.

## Acknowledgments

We thank Dr. Ben Parker for assistance with aphid symbiont infections, fungal infection protocols, and thoughtful feedback. We thank Manasa Peddineni for assistance with experiments. We thank Gerardo lab members for providing helpful comments on the manuscript and Tiger Li for comments on statistical analyses.

## Author contributions

WHT designed and carried out experiments, and wrote the manuscript. MR, KH designed and carried out experiments, and edited the manuscript. NMG designed experiments and edited the manuscript. TA, FL, and JB assisted with experiments and provided comments on the manuscript.

